# Dynamics of brain-fluid circulation are altered in the mature-onset Tet-off APP mouse model of amyloidosis

**DOI:** 10.1101/2022.03.11.483807

**Authors:** Inès R.H. Ben-Nejma, Aneta J. Keliris, Verdi Vanreusel, Peter Ponsaerts, Annemie Van der Linden, Georgios A. Keliris

**Affiliations:** Bio-Imaging Lab, University of Antwerp, Universiteitsplein 1, 2610 Wilrijk, Antwerp, Belgium; Laboratory of Experimental Hematology, Vaccine and Infectious Disease Institute (Vaxinfectio), University of Antwerp, Universiteitsplein 1, 2610 Wilrijk, Antwerp, Belgium; Research in dosimetric applications, SCK CEN, Boeretang 200, 2400 Mol, Antwerp, Belgium

**Keywords:** glymphatic system, mature-onset Tet-off mice, forebrain amyloidosis, amyloid-beta, DCE-MRI, inflammation

## Abstract

Alzheimer’s disease (AD), the most common type of dementia, is an incurable brain disorder characterised by the progressive build-up of toxic amyloid-beta (Aβ) and tau protein aggregates. AD gradually inflicts cognitive functions of an individual such as memory, thinking, reasoning, and language by degrading synaptic function and the integrity of neuronal networks. It has been recently suggested that the efficacy of different brain-clearance systems like the glymphatic system (GS), involved in the removal of toxic waste and homeostatic balance, plays a key role in the pathology of AD. Moreover, the observed coupling between brain fluid movement and global brain activity implies that an alteration of the neuronal network integrity can impact brain fluid circulation throughout the brain and thereby the efficacy of the GS. Here, we investigated the dynamics of brain fluid circulation in Tet-Off APP (AD) mice, a mature-onset model of amyloidosis in which we have recently shown a deterioration of neuronal network integrity by resting-state fMRI. By utilizing dynamic contrast enhanced-MRI and gadoteric acid (Gd-DOTA) T_1_ contrast agent injected into the cisterna magna, we demonstrated that brain fluid exchange was significantly altered in 14-month-old AD mice compared to control littermates. More specifically, AD mice showed higher Gd-DOTA accumulation in areas proximal to the injection cite and computational modeling of time courses demonstrated significantly lower inflow time constants relative to the controls. Immunohistochemistry demonstrated abundant amyloid plaque burden in the forebrain of the AD group coinciding with extensive astrogliosis and microgliosis. The neuroinflammatory responses were also found in plaque-devoid regions, potentially impacting brain fluid circulation.

## INTRODUCTION

Alzheimer’s disease (AD) is a progressive and still incurable devastating neurodegenerative disorder, clinically identifiable by a gradual cognitive decline, eventually leading to dementia and death (Dubois et al. 2016; Long and Holtzman 2019). AD is primary characterised by the accumulation of amyloid-beta (Aβ) and tau abnormal proteins deposits, which coincides with the disruption of the neuro-glial-vascular unit function and inflammation (Querfurth and LaFerla 2010; Zlokovic 2011; Soto-Rojas et al. 2021). The efficacy of debris removal via interacting brain clearance systems is believed to play an important role in the pathology of AD (Zlokovic et al. 2000; Weller et al. 2008; Iliff et al. 2012; Aspelund et al. 2015; Tarasoff-Conway et al. 2015; Louveau et al. 2017; Hladky and Barrand 2018; Mestre, Mori, and Nedergaard 2020). The more recently discovered glymphatic and brain lymphatic systems drew particularly attention and became of keen interest in the scientific community. The glymphatic system (GS), a brain-wide perivascular network of cerebrospinal fluid (CSF) and interstitial (ISF) fluid exchange that is facilitated by aquaporin-4 (AQP4) water channels expressed at astrocytic end-feet, has been shown to be critically involved in the clearance of Aβ and tau proteins from the brain (Iliff et al. 2012; Xie et al. 2013; Peng et al. 2016; Iliff et al. 2014; Harrison et al. 2020). Further, the CSF movement from the subarachnoid space to the paravascular space was shown to be driven by a combination of factors, such as cardiac and respiratory pulsations, sleep, vasomotion and CSF pressure gradients (Iliff, Lee, et al. 2013; Jessen et al. 2015; Xie et al. 2013; Iliff, Wang, et al. 2013; Mestre, Tithof, et al. 2018).

Importantly, the efficacy of glymphatic exchange was shown to decrease rapidly upon ageing in wild-type mice (Kress et al. 2014). Since ageing is the major risk factor for the prevalent in humans (~98%) late-onset AD (LOAD), age-related impairments of glymphatic circulation could be playing a key role in the progression of AD pathology. However, most of the commonly used transgenic models in AD research were generated based on genetic autosomal dominant mutations of early-onset AD (EOAD; <65 years old) (Jankowsky and Zheng 2017), such as the APP/PS1 and the 5xFAD (Radde et al. 2006; Oblak et al. 2021). One potential drawback these transgenic models have in common is the transgenic gene overexpression or overproduction of amyloid/tau during critical postnatal brain development phases.

Following the discovery of the GS and its implicated importance in the pathology of AD, several groups have used EOAD models to study glymphatic clearance and the factors governing its efficacy in AD. In effect, it was shown that glymphatic transport was affected in both young and old APP/PS1 mice compared to wild-type littermates (Peng et al. 2016). Further, Xu et al. showed that deletion of AQP4 in APP/PS1 mice resulted in aggravation of amyloid-β accumulation and memory impairment (Xu et al. 2015; Feng et al. 2020). In addition, the perivascular localisation of AQP4 channels is postulated to be essential in maintaining the efficacy of the GS and has been shown to be declining with age (Kress et al. 2014), but to be also related to certain stages of AD pathology (Zeppenfeld et al. 2017; Yang et al. 2011).

On the other hand, ultra-fast magnetic resonance encephalography (MREG) imaging at rest demonstrated unique spatiotemporal patterns of low frequency signals (<0.01 Hz), which were associated with glymphatic dynamics of CSF movement in the human brain (Kiviniemi et al. 2016). In addition, it has been demonstrated that during non-rapid eye movement (NREM) sleep, CSF dynamics are coupled with global resting-state functional MRI (rsfMRI) signals, implying a neural origin (Fultz et al. 2019). Moreover, Han and colleagues pointed out the strong coupling between global rsfMRI signals and CSF flow, which was correlated with stages of AD-related pathology including the cortical Aβ levels, suggesting a potential link to glymphatic flow and brain waste clearance (Han et al. 2021). Thus, the brain activity-associated large-scale neuronal modulations may directly impact the efficacy of the glymphatic clearance in the brain parenchyma. Moreover, ageing was associated with reduced and more fragmented slow-wave sleep, particularly in AD (Lloret et al. 2020; Mander, Winer, and Walker 2017), but also with a decreased CSF flux, where a massive depolarization of astrocytic AQP4 was found (Kress et al. 2014).

Furthermore, the dysfunction of the resting-state neuronal networks detected at advanced stages of AD in humans (Badhwar et al. 2017; Greicius et al. 2004) was also observed in several different transgenic EOAD amyloidosis mouse models overexpressing amyloid precursor protein (APP) during the brain postnatal development (Grandjean et al. 2016; Shah et al. 2013). Recently, we have reported the impairment of global resting-state neuronal network integrity in a mature-onset Tet-Off APP mouse model of amyloidosis, in which the APP overexpression was ‘turned-on’ in adulthood (3-month-old mice) when the brain can be considered as mature. This important manipulation ensures that APP overexpression and Aβ overproduction does not occur during brain postnatal development that could create false-positive phenotypes unrelated to AD (Ben-Nejma et al. 2019). For instance, circulating toxic Aβ species might have a different impact on neuronal circuits, cell signalling or synapse formation in mature and immature mice (Jankowsky and Zheng 2017; Sri et al. 2019).

With an eye on the bidirectionality of interactions between CSF circulation and global rsfMRI signals, here we decided to investigate the efficacy and dynamics of CSF-ISF exchange in Tet-Off APP mice, in which we previously observed the dysfunction of global neuronal networks at stages of advanced amyloidosis. We sought to investigate if and how the dynamics of brain-fluid circulation could be affected in this model, in which a potential obstruction of the glymphatic pathways was to be expected in areas with heavy amyloid plaque load such as the forebrain, and importantly in comparison to aged littermates in order to differentiate changes related to ageing from those driven by AD pathology.

## MATERIALS AND METHODS

### Mouse strain, dox treatment and housing

Generation of the Tet-Off APP transgenic mice has been described in detail in our previous study (Ben-Nejma et al. 2019). Briefly, this inducible model of amyloidosis allows a time-controlled expression of a chimeric mouse/human APP695 transgene using the Tet-Off system. The bigenic tetO-APPswe/ind (line 107) animals (strain B6.Cg-Tg(tetO-APPSwInd)107Dbo/Mmjax, referred to as AD mice in the manuscript) were bred in-house by crossing APP mice, in which a tetracycline-responsive (tetO) promoter drives the expression of the chimeric APP transgene bearing the Swedish and Indiana mutations (mo/huAPP695swe/ind), with transgenic mice expressing the tetracycline-Transactivator (tTA) gene (strain B6;CBA-Tg(Camk2a-tTA)1Mmay/J). The single transgenic tTA and APP males (Prof. Dr JoAnne McLaurin, Sunnybrook Health Sciences Centre, Toronto, Canada) were initially crossed with non-transgenic females on a C57BL6/J background (Charles River, France) to establish the single transgenic colonies. Since the tTA transgene is under control of the CaMKIIα promoter, the bigenic mice express APP in a neuron-specific manner at moderate levels, essentially in the forebrain (Jankowsky et al. 2005). The APP expression was ‘turned-off’ up to the age of 3 months (3m, young adult mice) by feeding females with litters and weaned pups with a specific chow supplemented with doxycycline (DOX), a derivate of tetracycline (antibiotics, 100 mg/kg Doxycycline diet, Envigo RMS B.V., The Netherlands) from P3 up to 3m. To induce APP expression in the bigenic mice, all in-house bred transgene carrier and non-transgenic carrier (NTg) littermates were switched to a regular chow from 3m onward until the day of the surgery (14m old) resulting in a total APP expression duration of 11m. Of note, we reused the animals that previously completed a longitudinal rsfMRI study, presented in our recent work (Ben-Nejma et al. 2019). The mice genotypes are indicated in figure schemes or legends throughout the manuscript. Animals were housed in an environment with controlled temperature and humidity and on a 12h light–dark cycle, and water was provided ad libitum. All experiments were approved and performed in strict accordance with the European Directive 2010/63/EU on the protection of animals used for scientific purposes. The protocols were approved by the Committee on Animal Care and Use at the University of Antwerp, Belgium (permit number 2014–76).

### Surgery

The injection of gadolinium (Gd)-based T_1_ contrast agent, gadoteric acid (Gd-DOTA) into the cisterna magna (CM) was performed in spontaneously breathing mice anesthetized with 2% isoflurane (3% for short induction) delivered in oxygen by adapting a previously reported protocol (Gaberel et al. 2014; Xavier et al. 2018). Briefly, the animal was positioned in a custom-made stereotaxic frame with its head pointing down to expose the CM. A midline incision was made from the occipital crest down to the first vertebrae. Then, the underlying muscles were gently separated and maintained pulled aside using two curved forceps. The CM appeared as a small, inverted triangle overlaid with the translucent dural membrane, in between the cerebellum and the medulla. After exposing the CM, 2.5 μl of 50 mM solution of Gd-DOTA (DOTAREM®, Guerbert, France) was injected at 0.55 μl/min via a pulled haematological glass micropipette attached to a nanoinjector (Nanoject II Drummond). To avoid leakage, the micropipette was left in place for additional 5 minutes and the incision was closed with biocompatible superglue. The body temperature was maintained at 37.0°C with a heating pad.

### MRI acquisition

Following surgery, animals were positioned in MRI-compatible cradle/bed (animal in prone position) using MRI compatible mouse stereotactic device, including a nose cone to deliver anaesthetic gas 2% isoflurane (Isoflo^®^, Abbot Laboratories Ltd., Illinois, USA) administered in a gaseous mixture of 33% oxygen (200 cc/min) and 67% nitrogen (400 cc/min). During the MRI acquisition, the mice were allowed to breathe spontaneously. The physiological status of the animals was closely monitored during the entire acquisition. The respiration rate was maintained within normal physiological range (80-120) breaths/min using a small animal pressure sensitive pad (MR-compatible Small Animal Monitoring and Gating System, SA instruments, Inc.). The body temperature was monitored by a rectal probe and maintained at (37.0 ± 0.5) °C using a feedback controlled warm air system (MR-compatible Small Animal Heating System, SA Instruments, Inc.).

All imaging measurements were performed on a 9.4T Biospec MRI system (Bruker BioSpin, Germany) with the Paravision 6.0 software (www.bruker.com) using a Bruker coil set-up with a quadrature volume transmit coil and a 2×2 surface array mouse head receiver coil. Axial and sagittal 2D T2-weighted Turbo RARE images were acquired to ensure uniform slice positioning (RARE; TR/TE 2500/33 ms; 9 slices of 0.7 mm; FOV (20×20) mm^2^; pixel dimensions (0.078×0.078) mm^2^). Dynamic contrast-enhanced MRI (DCE-MRI) acquisition was performed using a 3D T1-weighted FLASH sequence (3D T1-FLASH; TR/TE 15/4.3 ms; flip angle 20°) in the sagittal plane. The field-of-view (FOV) was (18×15×12) mm^3^ and the matrix size (96×96× 64), resulting in voxel dimensions of (0.188 × 0.156× 0.188) mm^3^. The DCE-MRI scans were acquired every 5 min and started 30 min up to 150 min after the contrast agent injection. An overview of the experimental setup is summarized in Figure 1.

**Figure 1.**
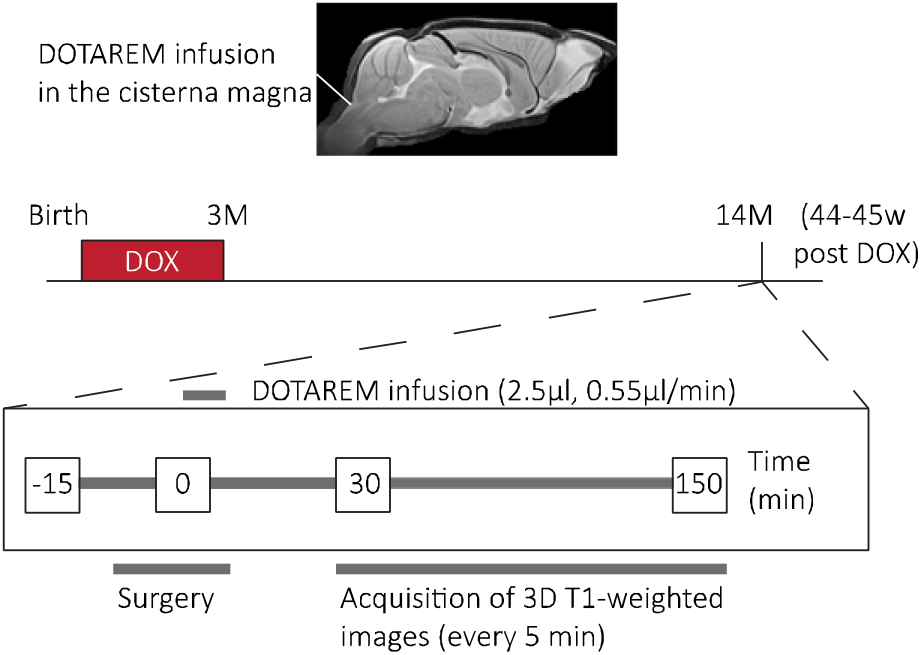
Overview of the experimental setup. The 14 m old animals, AD and CTL, underwent the surgery after 11 months of amyloid-beta expression. The surgery started a few minutes after the anaesthesia induction (time point=-15 min) and the start of continuous Gd-DOTA infusion (2.5 μl, 0.55 μl/min) into the cisterna magna refers to the time point t = 0. The DCE-MRI acquisition (120 min) started 30-min post Gd-DOTA injection, every 5-min, up to 150-min post injection.

### MRI data pre-processing

Pre-processing of the DCE-MRI data was performed using Advanced Normalisation Tools (ANTs) including realignment, spatial normalisation and creation of a 3D study template. First, a mean image has been created across the time series for each subject and a mask larger than the brain has been delineated on it using AMIRA 5.4. Then, this broad mask has been applied to the DCE-MRI images to remove the surrounding muscle tissue. In parallel, a study-specific 3D-template based on the last scan of the non-transgene carrier (NTg) group was created using a global 12-parameter affine transformation followed by a nonlinear deformation protocol. This template was used to estimate the spatial normalisation parameters of the mean images. Next, the realignment parameters of all masked DCE-MRI images within each session to the masked mean image were first estimated, using a symmetric image normalisation method (SyN transformation). Then, the transformation parameters of the realignment and the spatial normalisation were applied to the DCE-MRI images in one resampling step.

Signal intensity normalization was performed in MATLAB (MATLAB R2020a, The MathWorks Inc. Natick, MA, USA). First, an ellipsoid-shaped region-of-interest (ROI) of 141 voxels was delineated in a cortical area where the variability of the intensity over time was negligible for the baseline and saline groups. More specifically, a time frame of six consecutive scans was selected based on the least changes in the time traces for this specific mask (i.e., intensity values remained approximately constant). Therefore, the mean intensity value of this mask for these six consecutives scans was used to convert each voxel of all images to percent signal change. Finally, a smoothing step was performed with a 3D Gaussian kernel of radius twice the voxel size. A second mask restricted to the brain was applied to all images.

### MRI data analysis

Five groups of 14-month-old mice (14 m) were subjected to DCE-MRI experiments: a non-transgene carrier (NTg) non-injected group (NONE, N=3), a NTg saline-injected group (SAL, N=3), a NTg Gd-DOTA injected group (CTL_1_; N=5), a tTA Gd-DOTA injected group (CTL_2_; N=3) and a bigenic Tet-Off APP Gd-DOTA injected group (AD; N=7). In total 21 mice (14 m, mixed in gender) were scanned that were reared on Dox diet until 3m of age. Five animals (2 NTg, 1 tTA and 2 AD) have been removed due to surgery failure. Given that tTA animals (CTL_2_) do not produce sAβ or amyloid plaques and showed no difference to the NTg littermates (CTL_1_) in our previous resting state experiments (Jankowsky et al. 2005; Ben-Nejma et al. 2019), these two groups of animals were combined and are further referred to as control group (CTL, N_CTL_ = 5).

First, principal component analysis (PCA) was performed per group on the average of the spatially smoothed and normalised time-courses. As more than 99% of the data variability could be explained by the three largest components, we used them to reconstruct PCA-based time courses that effectively reduced high frequency noise from the data. Subsequently, a hierarchical clustering (ward linkage, maximum 15 clusters) was performed on the reconstructed times courses of all animal groups. This analysis allowed for the identification of clusters of voxels with similar time courses and to observe the patterns across groups. Then, to have a fair comparison in the same voxels, the clusters of either the CTL group or the AD group were used to compare the voxel averaged time-courses of the CTL and AD groups and to assess the dynamics of glymphatic flow based on modelling. To this end, we sub-selected the clusters with at least 100 voxels that showed a difference between maximum and minimum intensity of more than 10%. This criterion was selected based on the variability observed in the NONE and SAL groups for which the time-courses were flat as expected (see results). Subsequently, PCA was also performed on a subject-by-subject basis to allow for statistical analysis between the groups (CTL vs. AD). Analyses were performed: a) on six predefined hypothesis driven regions of interest (ROIs), and b) on the clusters defined based on the group level PCA of the CTL group as described above. For ROI-based analysis, six relevant ROIs (olfactory bulb, hippocampus, medulla, pons, aqueduct, and cerebellum) were delineated with MRIcroGL based on the Paxinos atlas (3^rd^ edition). Then, for each subject, the mean of the PCA-based time-courses over the voxels included in each ROI was extracted and the area under the curve (AUC) was calculated and used for statistical analysis (t-test across the two groups). For cluster-based analysis, the mean over the voxels included in each cluster was extracted for each subject separately and then the time-courses of each cluster were fitted using a model with two exponentials based on the following formula: 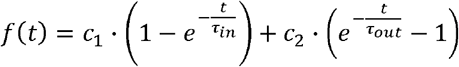, with *c*_1_ and *c*_2_ representing gain constants, τ_in_ the inflow time constant, and τ_out_ the outflow time constant. The estimated τ’s for the inflow and outflow for each cluster per subject were then used for statistical comparison (t-test) across the two groups.

### Statistical analyses

For the ROI-based analysis, a two-sample *t*-test was performed on the area under the curve (AUC) values per ROI for the CTL vs. AD groups using MATLAB (MATLAB R2020a). Similarly, for cluster-based analysis, a two-sample *t*-test was performed on the τ_in_ and τ_out_ values per cluster across the two groups. Significance was defined with a criterion α=0.05. All results are shown as mean ± standard errors.

### Immunohistochemistry

Brain samples were collected directly after the MRI acquisition (N_NTg_=3; N_AD_=3) as described previously (Ben-Nejma et al. 2019). Briefly, the mice were deeply anesthetized with an intraperitoneal injection of 60 mg/ kg/BW pentobarbital (Nembutal; Ceva Sante Animale, Brussels, Belgium), followed by a transcardial perfusion with ice-cold PBS, and with 4% paraformaldehyde (Merck Millipore, Merck KGaA, Darmstadt, Germany). Brain-samples were afterwards surgically removed and post-fixed in 4% paraformaldehyde for 4h. Next, the fixed brains were freeze-protected using a sucrose gradient (sucrose, Sigma Aldrich): 2h at 5%, 2h at 10%, and overnight at 20%. Then, the brain samples were snap frozen in liquid nitrogen and stored at −80°C. Finally, 14-μm-thickness sagittal brains sections were cut using a cryostat (CryoStar NX70; ThermoScientific).

For immunofluorescence analyses, the following primary antibodies were used: chicken anti-GFAP (Abcam ab4674, 1:1000), rabbit anti IBA-1 (Wako 019-19741, 1:1000), rabbit anti-AQP4 (Sigma-Aldrich HPA014784, 1:100) and the following secondary antibodies: donkey anti chicken (Jackson 703-166-155, 1:1000), goat anti rabbit (Jackson 111-096-114, 1:1000) and donkey anti rabbit (Abcam AF555, 1:1000). Moreover, the amyloid plaques were stained with Thioflavin-S (Santa Cruz Biotechnology, sc-215969) and the vessels with lectin (Labconsult VEC.DL-1174 (green) or VEC.DL-1177 (red), 1:200). After the staining, the sections were mounted using Prolong Gold Antifade (P36930; Invitrogen).

Immunofluorescence images of GFAP/lectin, Iba1/lectin, AQP4/lectin and Thioflavin-S/lectin stainings were acquired using an Olympus BX51 fluorescence microscope equipped with an Olympus DP71 digital camera and the image acquisition was done with CellSens Imaging Software (Olympus, Tokyo, Japan, http://www.olympus-global.com). Obtained images were visually evaluated by at least three co-workers to ensure the selection of representative images. The 3 edition of the mouse brain atlas from Paxinos and Franklin was used as reference for the localization of the regions of interest. Images were further processed with ImageJ Software 1.52k (National Institutes of Health) and artificially pseudo-coloured in the representative images.

## RESULTS

### Differences in the spatiotemporal distribution of Gd-DOTA in the Tet-Off APP mice vs. controls

We first investigated the efficacy of glymphatic transport in 14m old Tet-Off APP (AD) and control (CTL) mice by assessing the dynamics and distribution patterns of contrast agent (Gd-DOTA) in the brain. For this, T1-weighted contrast-enhanced MR images were acquired sequentially from 30 min to 150 min following the labelling of CSF upon infusion of Gd-DOTA into cisterna magna (t=0). We detected clear differences in the dynamics of brain-wide distribution patterns of Gd-DOTA, with reduced rostral glymphatic flow in the AD group and higher accumulation of contrast agent within caudal regions of the brain (see Figure 2C for an overview of the Gd-DOTA distributions in the brains of the two groups of animals).

**Figure 2.**
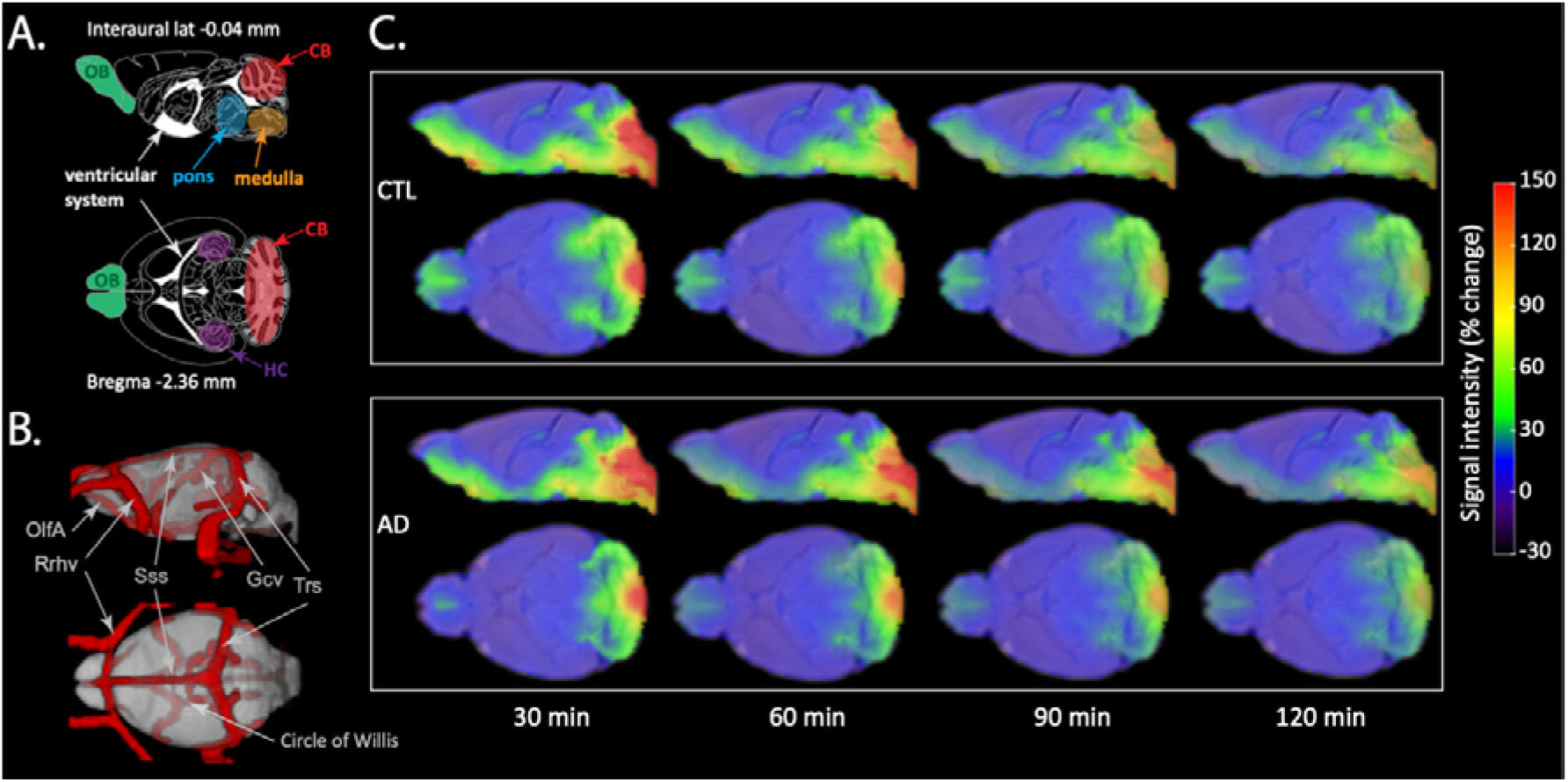
Spatial distribution maps of Gd-DOTA in the AD and CTL mice over time. A. Schematic representation of the sagittal plane (interaural lateral −0.04 mm) and transverse plane (Bregma level - 2.36 mm) from the Paxinos atlas with highlighted key regions such as olfactory bulb (OB), cerebellum (CB), hippocampus (HC), pons, medulla and ventricular system. B. 3D visualisation of major blood vessels based on a recent vascular brain mouse atlas that was aligned to the template of this study (vascular atlas from (Hinz et al. 2021)). The depicted vessels including OlfA (olfactory artery), Rrhv (rostral rhinal vein), Gcv (cerebral vein of Galen), Sss (superior sagittal sinus), Trs (transverse sinus) are located in the vicinity of regions with enhanced contrast. C. The spatial distribution of the MRI contrast agent in the CTL (top panel) and AD mice (bottom panel) at 30-min, 60-min, 90-min and 120-min post intracisternal infusion shown in sagittal and transverse views. The colour scale represents the average percent signal intensity change, with dark blue/red indicating low/high percent signal intensity change, respectively.

In line with the literature, characteristic patterns of GS-related contrast agent distribution were observed in our CTL mice, with contrast enhancement of the paravascular routes and the adjacent parenchyma (Iliff et al. 2012; Iliff, Lee, et al. 2013; Gaberel et al. 2014; Gakuba et al. 2018). More specifically, the CTL mice showed high accumulation of Gd-DOTA at 30-min post injection, reflected as high signal intensity, in the cisterna magna, ventricular system, ventrally along the circle of Willis and olfactory paravascular path (Figure 2A-C, top panel), as well as in caudal parenchymal brain regions including the brainstem (i.e., medulla and pons). Other regions of detected contrast enhancement included the cerebellum, the pituitary recess, the ventral part of the thalamus and the olfactory bulb. In addition, contrast enhancement was also observed in the areas adjacent to the transverse sinuses (see also Additional Figure 1). Starting from 60-min onward, the signal intensity of the Gd-DOTA started to fade drastically, except for the regions located in the vicinity of the cisterna magna, which showed higher contrast compared to the rest of the brain.

At the imaging onset (t=30 min), the Gd-DOTA distribution for the AD group was spatially largely similar to the controls, while differences became more pronounced over time (Figure 2C, bottom panel). In effect, at 30-min post-infusion, we noticed that the T_1_ signal intensity was higher at the caudal brain regions, including the ventral part of the cerebellum, medulla, pons, while lower contrast enhancement was observed within the olfactory bulb. From 60-min post-infusion onward, the heterogenicity in spatial differences was evident, with a higher signal intensity of the Gd-DOTA, retained for a longer time in the brainstem (i.e., pons and medulla) and the ventral part of the cerebellum (CB). Compared to the CTL group, a lower contrast enhancement was detected within the olfactory bulb.

### Caudal retention and reduced rostral flow of Gd-DOTA in AD mice

To investigate the glymphatic CSF-ISF exchange in more detail, the quantification of Gd-DOTA accumulation was assessed within specific anatomical regions. To this end, a ROI-based analysis was performed with the outcome being illustrated in Figure 3. In line with results highlighted in the previous section, the AD mice showed significant differences in signal intensity, with much higher and slower accumulation, time-to-peak, as well as longer retention in the medulla (p = 0,0278, Figure 3D) and the pons (p = 0,0191, Figure 3E) compared to CTL mice, while no differences were found in the olfactory bulb, cerebellum and aqueduct (Figure 3A-C, F). Further, a trend for slightly higher accumulation of Gd-DOTA, albeit not statistically significant, within hippocampus regions (Fig 3B, p = 0,1159) was found for CTL littermates. Thus, these data indicated that the glymphatic pathways of CSF circulation were altered in AD mice, and transport of Gd-DOTA from the cisterna magna towards the rostral and dorsal aspects of the brain were reduced and accompanied by prolonged accumulation at the caudal regions (medulla and pons) that are free of amyloid plaques (Jankowsky et al. 2005).

**Figure 3.**
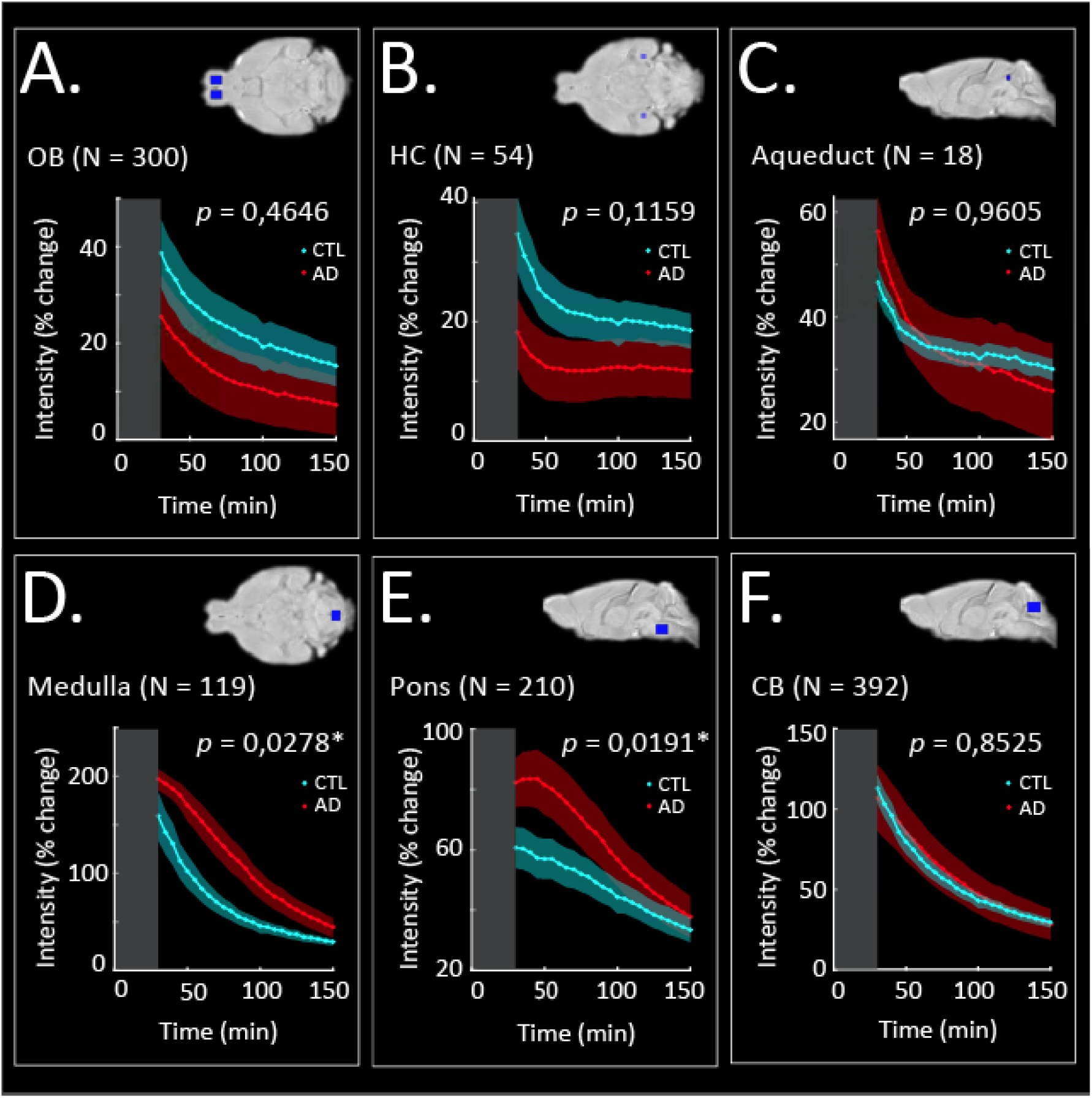
Region-of-interest (ROI)-based analysis. Average signal intensity changes over time reflecting Gd-DOTA contrast agent distribution in the OB (A), the HC (B), the aqueduct (C), the medulla (D), the pons (E) and the CB (F) for the CTL (cyan) and the AD (red) groups of mice. The grey rectangle in each panel represents the pre-acquisition time (acquisition started at 30-min post Gd-DOTA infusion) with zero indicating the time of injection. An overview of the brain location of each ROI can be found in the top right corner of each panel (blue rectangles). OB, olfactory bulb; HC, hippocampus; CB, cerebellum; N, ROI size in voxels, *; significant difference in the area-under the curve (AUC) across the two groups indicates a differential distribution of contrast agent (p<0.05; see Materials and Methods).

### Clustering of voxel time-courses reveals differential Gd-DOTA spatial patterns in AD and CTL mice

Given the observed differences in the distribution of Gd-DOTA in the selected ROIs over time (Figures 2 & 3), we sought to evaluate whether the similarity of the time-courses of glymphatic circulation across the brain, could provide information about the distribution patterns and pathways. To this end, we performed cluster analysis on the group average PCA based time-courses by using the three largest principal components that could explain ~99% of the data variability. The resulting clustering maps for each group are shown in Figure 4A and the corresponding time-courses in Figure 4B.

**Figure 4.**
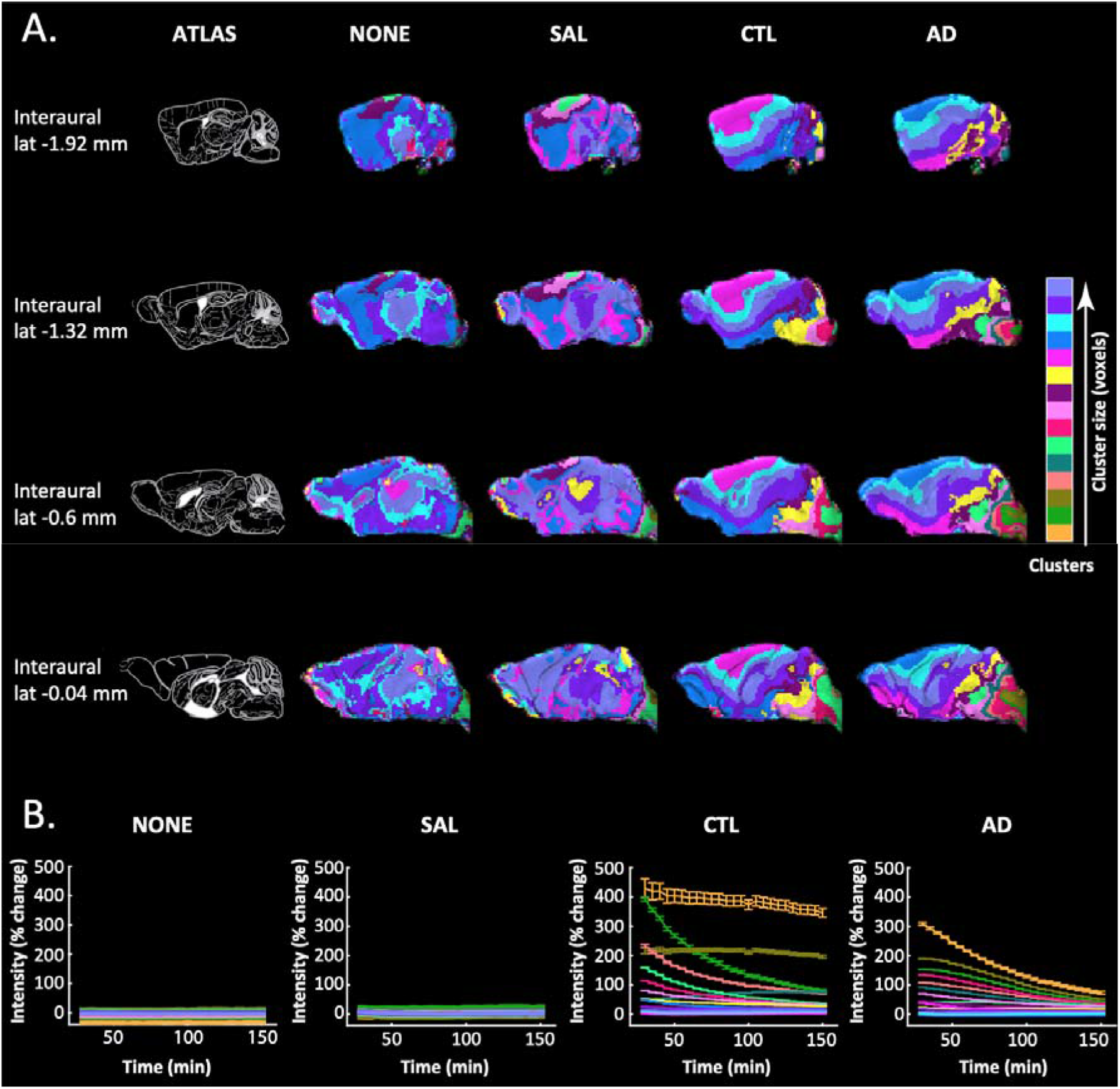
Clustering of MRI voxel time-courses reflecting Gd-DOTA contrast agent distribution over time in the Tet-Off APP mouse model (AD) and three control groups: non-injected (NONE), saline-injected (SAL), and Gd-DOTA injected (CTL). A. On the leftmost column (ATLAS), sagittal planes from the Paxinos atlas at different interaural locations indicate four selected, representative brain slices. In other columns, the visualization of 15 clusters for each group (each cluster shown with a different colour) demonstrates brain areas that share similar distributions of contrast agent. Colours on the colour-bar indicate the sorting of clusters (in number of voxels) from smallest to largest in each group but note that same colours in different groups do not entail the same sizes and that it is not a linear scale but rather a sorted legend. B. The voxel-average time-courses (mean ± SD) of each cluster are shown for each group in the same colour as the respective cluster in A. Note that non-injected groups (NONE, SAL) show flat distributions of signal intensity over time that simply relate to slight signal amplitude differences without contrast agent contribution. On the other hand, injected groups demonstrated large increases in signal intensity starting at 30 min post-injection of Gd-DOTA (start of MRI acquisition) declining over time in most of the clusters (clearance of contrast agent) but also note the accumulation of contrast agent in a couple clusters.

As expected by the absence of contrast agent, the clustering maps of the non-injected (NONE) and saline-injected (SAL) groups did not show a particular structure, but rather consisted of some very large homogeneous clusters and some smaller ones distributed randomly over the brain (Figure 4A). Moreover, as shown in Figure 4B, the time-courses for the SAL and NONE groups were approximately constant over time and close to 0% intensity change, essentially showing that the percent signal intensity change over time was minimal and much smaller in relation to the contrast injected groups. The small variability in the intensities of different clusters, approximately ±5%, was used to guide the criterion for selecting clusters for further analysis in the injected groups, i.e., the clusters showing substantial intensity changes over time and thus reflecting changes in the distribution of contrast agent rather than simple signal variability (see Materials and Methods).

On the contrary to the NONE and the SAL groups, the CTL and AD groups that received intracisternal Gd-DOTA injections, displayed distinct clustering patterns. We could clearly observe a gradual spreading of the clusters from the posterior of the brain proximal to the cisterna magna towards the anterior of the brain and the olfactory bulb and dorsally towards the cortex (Figure 4A). Further, we noticed that for the AD group, the clusters proximal to the cisterna magna were more heterogeneous (fragmented) compared to the CTL. In addition, the time-courses of the clusters in the AD mice showed a delay in reaching the maximum peak compared to those in the CTL (Figure 4B).

### The dynamics of CSF-ISF exchange are altered in the AD mice

Based on the obtained clustering maps and time curves, we sought in depth comparison between the AD and CTL mice to unravel differences in the dynamics and kinetics of CSF-ISF exchange. To this end, we selected clusters of interest from the CTL and AD groups based on a minimum size (in number of voxels) and peak intensity changes (see Materials and Methods section for selection criteria) and extracted their average time-courses (Figure 5A-D). Next, we extracted the time-courses from the same clusters in each animal and fitted them using a double exponential model with parameters representing the time constants for the inflow (τ_*in*_) and the outflow (τ_*out*_) of the Gd-DOTA. The group average time-courses and the corresponding fitting curves using the clusters from the CTL and AD groups are shown in Figure 5E (N=6) and 5F (N=10), respectively.

**Figure 5.**
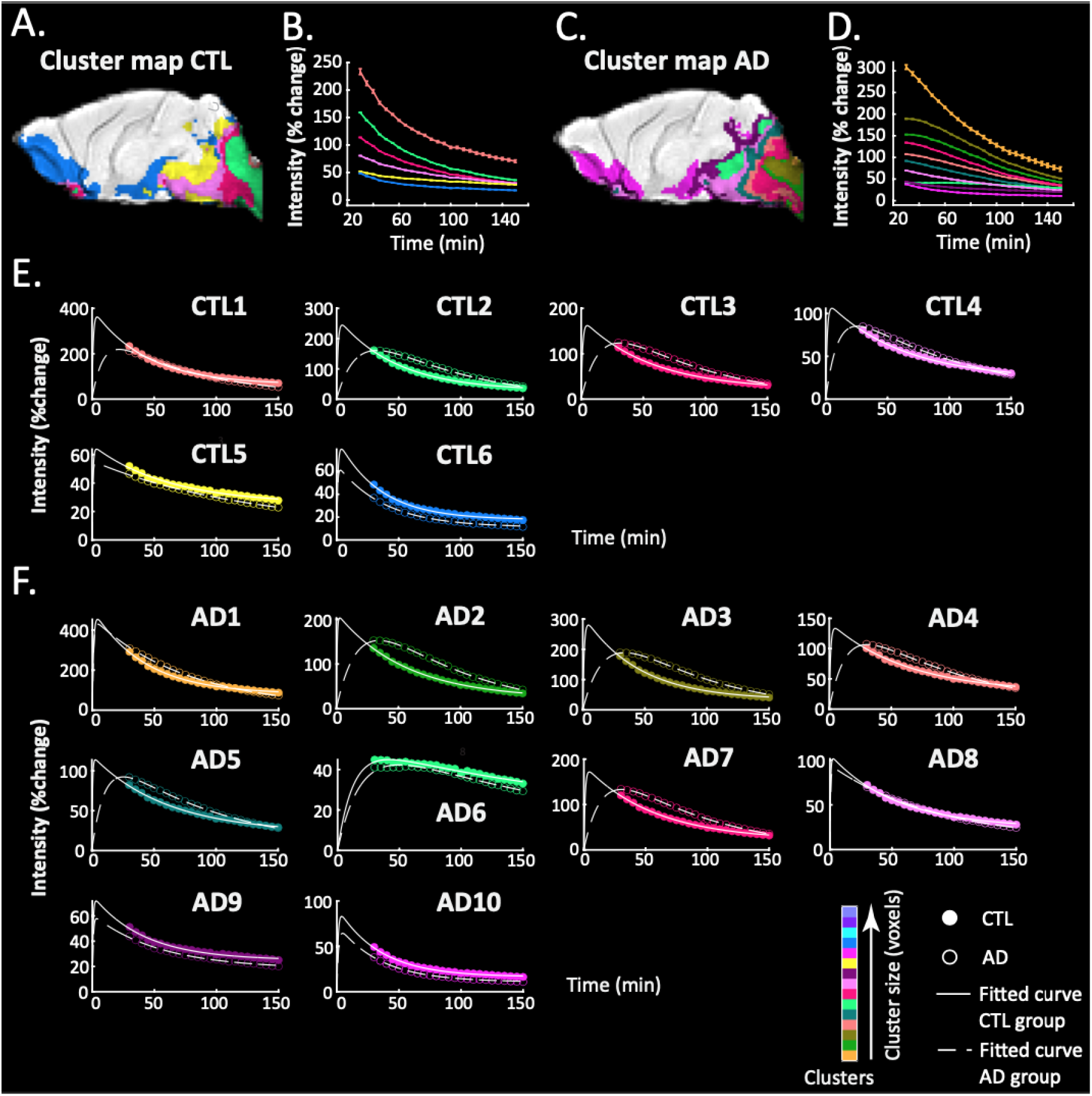
Model-fitting of cluster time-courses. (A, C) Clusters of the CTL and AD groups respectively selected for further analysis based on a minimum signal-intensity change and minimum number of voxels criterion (see Materials and Methods). (B, D) Time-courses of the selected clusters in the CTL and AD group respectively. (E) Model-fitting of the time-courses of the selected clusters from the CTL group (solid lines, filled circles) is compared to model-fitting of the same cluster of voxels in the AD group (dashed lines, open circles). (F) Similarly, model-fitting of the selected clusters of the AD group (dashed lines, open circles) is compared to model-fitting of the same cluster of voxels in the CTL group (solid lines, closed circles). In each panel of E and F, lines demonstrate the model fit and circles the data points (also shown in B, D) for each cluster. Colours represent size sorting of the clusters based on the number of voxels (same as Fig. 4).

For the CTL based clusters (Fig. 5E), we noticed that those located in the vicinity of the injection site showed different profiles of data and fitted curves across the two groups (Fig. 5E; CTL1-4). While the CTL group curves were from the start of acquisition in the descending phase of signal intensity (representing the outflow of the Gd-DOTA), initial increases of signal intensity (representing inflow and accumulation of Gd-DOTA) were still occurring in the AD mice with signals reaching the maximum peak at later time points. Furthermore, the infiltration of Gd-DOTA into the brain parenchyma was significantly longer in the AD group. These observations were also consistent with the model fitted τ_in_ time constants representing contrast agent inflow, which showed significant differences across the two groups (see τ_in_ values of clusters CTL1-4 in Table 1). In contrast, two other clusters (CTL5 and CTL6), displayed similar patterns in both groups and no significant differences in the inflow were observed. Notably, in the case of the outflow time constant τ_out_, we found no significant differences between the AD mice and their CTL littermates for any of the CTL based clusters (Table 1).

**Table 1.** p-values of the two-sample t-test performed on the time constants τ_in_ and τ_out_ representing the inflow and outflow of DOTAREM for the CTL-based clusters. *, (p<0.05).

| *Cluster* | CTL1 | CTL2 | CTL3 | CTL4 | CTL5 | CTL6 |
| --- | --- | --- | --- | --- | --- | --- |
| $\tau_{in}$ | 0.0149* | 0.0248* | 0.0199* | 0.0274* | 0.1893 | 0.6197 |
| $\tau_{out}$ | 0.9047 | 0.2088 | 0.4702 | 0.9026 | 0.0955 | 0.6765 |

For the AD based clusters (Fig. 5F), even though their spatial distribution was different with more clusters in total, a similar effect - namely the inflow of Gd-DOTA was significantly slower in the AD mice - was observed particularly for those clusters in the vicinity of the injection site (Fig. 5F; AD2,3,4 & 7). This was also reflected in the statistics of the τ_in_ and τ_out_ time constants across the two groups (Table 2). Note that AD5, which consisted of voxels surrounding the aforementioned clusters and thus was receiving Gd-DOTA from them, also showed a statistical trend (p = 0.0563) for a difference in the inflow τ_in_ parameter across the two groups. In analogy to the CTL clusters, the clearance τ_out_ of Gd-DOTA was similar between both groups with no significant differences in any cluster (see p-values of τ_out_ in Table 2).

**Table 2.** p-values of the two-sample t-test performed on the time constants (τ) for the inflow and the outflow of the DOTAREM for the AD clusters. *, (p<0.05), ^#^, trend for statistical significance.

| *Cluster* | AD1 | AD2 | AD3 | AD4 | AD5 | AD6 | AD7 | AD8 | AD9 | AD10 |
| --- | --- | --- | --- | --- | --- | --- | --- | --- | --- | --- |
| $\tau_{in}$ | 0,2868 | 0,0209* | 0,0204* | 0,0327* | 0,0563<sup>#</sup> | 0,4011 | 0,0260* | 0,1161 | 0,1541 | 0,6050 |
| $\tau_{out}$ | 0,5503 | 0,7787 | 0,2947 | 0,7097 | 0,4027 | 0,3415 | 0,8662 | 0,2634 | 0,1631 | 0,9691 |

### Altered glymphatic transport in AD mice is associated with brain pathology at the microscale

To further clarify the observed differences in the dynamics of brain-fluid circulation in the AD group, we have also performed an ex vivo histological assessment to assess microscale alterations in multiple brain regions. To this end, we evaluated both the amyloid plaque burden (Thioflavin-S staining) and the inflammatory responses, such as astrogliosis (GFAP staining) and microgliosis (Iba1 staining) (Figure 6A-C). Furthermore, we evaluated AQP4 colocalization with the blood vessels (Lectin staining) (Figure 6D). A large amyloid plaque burden was observed throughout the brain of AD mice, including hippocampus, cortex, olfactory bulb and amygdala, with no deposits present in the cerebellum and the brainstem (Figure 6A), thereby consistent with the previously reported forebrain parenchymal amyloidosis for this Tet-off APP mouse model (Jankowsky et al. 2005). In terms of inflammatory processes, extensive astrogliosis and microgliosis were clearly present in the brain of AD mice, with reactive microglia and astrocytes greatly surrounding the vicinity of dense-core amyloid plaques (Figure 6B-C). Notably, astrogliosis and microgliosis were also observed in brain regions devoid of plaques, suggesting the spread of inflammatory signalling throughout the whole brain of the AD mice. Despite extensive astrogliosis, vessel density was not altered and AQP4 seemed to remain colocalised along the blood vessels in AD mice, regardless of the brain areas (Figure 6D).

**Figure 6.**
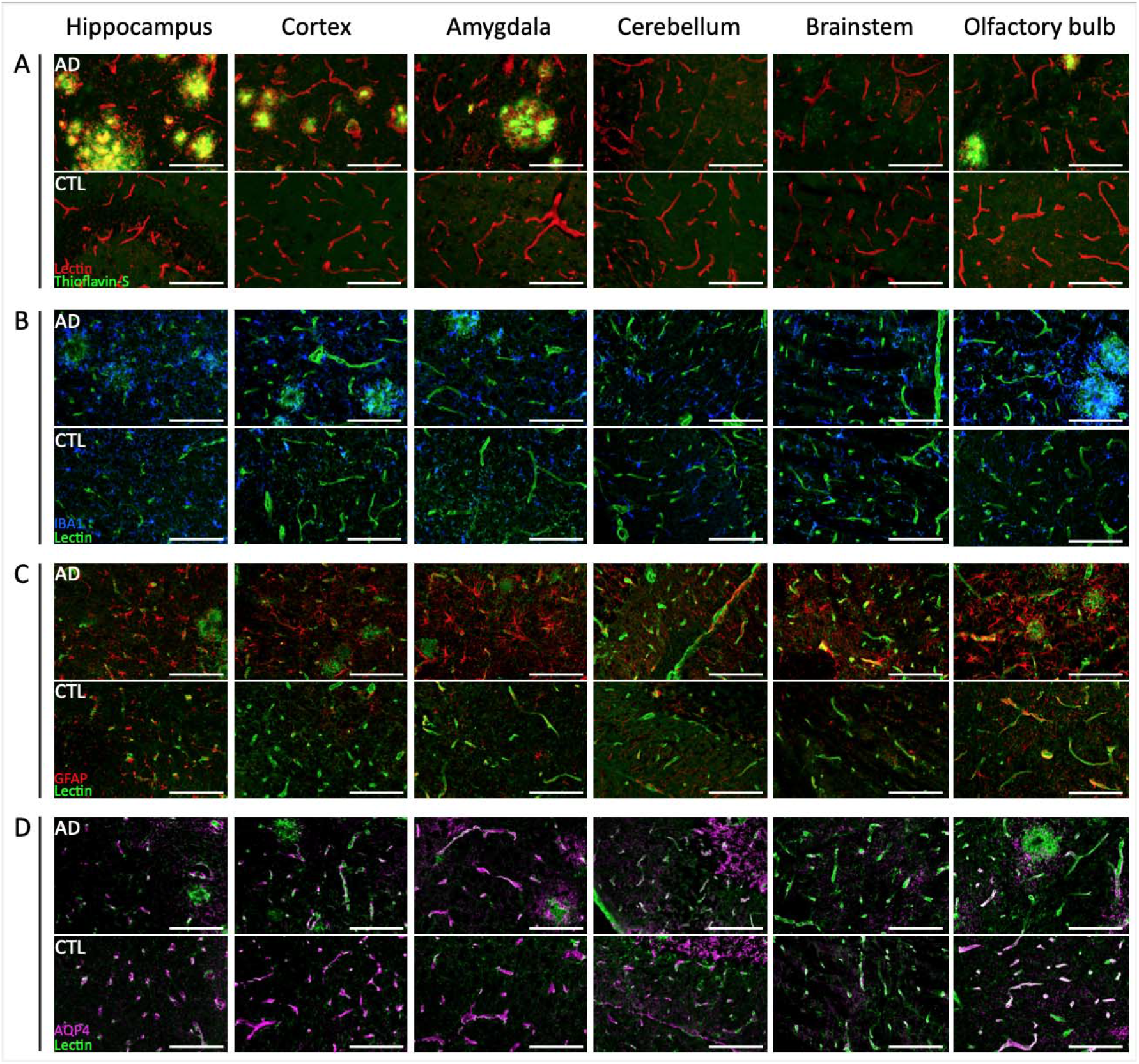
Representative images of ex vivo evaluation of amyloid plaque load, inflammatory responses and AQP4 in the Tet-Off APP (AD) and control (CTL) mice within key brain regions hippocampus, cortex, amygdala, olfactory bulb, cerebellum and brainstem. A. Amyloid plaques (green) stained with Thioflavin-S were detected in brain regions characteristic of the model, such as the hippocampus, the cortical areas and olfactory bulb. The vessels were labelled with lectin (red). B. Obvious sign of microgliosis (Iba1, blue) was observed surrounding the amyloid plaques and the surrounding tissue in the AD mice but not surrounding the vessels (lectin, green). C. AD mouse showed an accumulation of astrocytes (GFAP, red) surrounding the plaques, as well as the surrounding tissue and vessels (lectin, green) compared to control littermates, even in brain regions devoid of plaques (i.e., brainstem). D. AQP4 (magenta) is colocalised along the vessels (lectin, green) and its distribution does not seem to be different between both groups. Scale bar, 100 μm.

## DISCUSSION

Driven by recent findings linking AD and ageing to CSF circulation and neuronal network dysfunction, we performed this DCE-MRI study in a mature-onset mouse model of amyloidosis mimicking late-onset AD (LOAD) in humans. Particularly, to avoid possible confounding factors caused by APP overexpression during brain postnatal development, we used 14-month-old Tet-Off APP (AD) mice in which APP overexpression, and thus transgenic Aβ overproduction, was ‘turned-on’ at 3 months of age (‘mature’ adult) and compared their whole brain fluid circulation with non-plaque bearing control littermates.

We could demonstrate that AD mice and controls manifested clear and interesting divergence in their CSF-ISF exchange as captured by in vivo DCE-MRI (Figure 2, Figure 4). More specifically, we elucidated that the glymphatic transport of the Gd-DOTA tracer was considerably changed in AD mice compared to the control group with changes indicating a reduced and redirected flow (Figure 2C). Notably, we observed that the AD group showed slower infiltration and time-to-peak, longer retention, and high accumulation of CSF tracer within caudal regions of the brain, such as the medulla and the pons (Figure 3D-E). This was even more evident while looking at the profiles of the time-courses for both groups (Figure 4), where for AD mice the clusters of regions placed in the proximity of the injection side presented a delay in reaching the peak intensity compared to the controls that were already in the descending phase. Moreover, the modification of Gd-DOTA circulation was reflected in the differences of the inflow time constant τ_in_ (see Tables 1 and 2) with evidently longer and slower parenchymal infiltration of CSF tracer for the AD group.

Recently, we reported that the same AD mice showed large deterioration of global functional networks as reflected in resting-state functional MRI measurements with reduced functional connectivity within and between regions of the default-mode network (DMN) at the post-plaque stage (Ben-Nejma et al. 2019). In a recent study in humans, Han and colleagues analysed multimodal data from the Alzheimer’s Disease Neuroimaging Initiative (ADNI) and showed that the coupling between the global fMRI signal and CSF influx is correlated with AD pathology (Han et al. 2021). Thus, taken together, our findings from both studies are consistent with the work of Han and colleagues pointing towards mutual interactions between CSF flow and global brain activity in AD.

Our immunohistochemistry results in the AD mice showed wide-spread amyloidosis throughout several regions including cerebral cortex, hippocampus, olfactory bulb and amygdala, while cerebellum and brainstem remained free of amyloid-β deposits. Altogether, we reasoned that the high abundance of amyloid-β plaques across the forebrain of AD mice might have led to an obstruction and hindered the normal routes of CSF circulation within the cranial cavity. Previous studies demonstrated that blockage of normal routes can redirect and/or impair the CSF bulk flow (Peng et al. 2016; Wang et al. 2017; Ma et al. 2019). To this end, Ma and colleagues reported that after cisterna magna injection in glioma-bearing mice the CSF tracer was rerouted to the spine and spinal lymphatic outflow pathways due to the blockage of cranial CSF outflow routes (Ma et al. 2019). Further, Wang and colleagues showed that CSF flow was halted in the hemisphere affected by multiple microinfarctions and was slowed on the contralateral side (Wang et al. 2017). Also, in the context of amyloidosis models mimicking early-onset AD (EOAD) in humans, the glymphatic transport was greatly reduced in old APP/PS1 mice (12-18 m) (Peng et al. 2016). Note that in those mice, deposition of plaques begins at six weeks of age with amyloid-β spreading into several regions including cortex, hippocampus, striatum, thalamus, cerebellum, brainstem and limitedly vessels (Radde et al. 2006). Peng and colleagues also showed a deleterious effect of circulating soluble Aβ, which when injected into wild-type mice suppressed the influx of CSF tracer into the cortex (Peng et al. 2016).

Consistent with our results, altered glymphatic transport was also observed in humans. It was shown that glymphatic clearance in AD patients was reduced (de Leon et al. 2017) and delayed compared to healthy controls, demonstrated by a higher signal retention (Ringstad et al. 2018). Further, cognitively affected patients with idiopathic normal pressure hydrocephalus also displayed a delayed clearance of CSF tracer from entorhinal cortex (Eide and Ringstad 2019).

Importantly, glymphatic system decline was also found upon normal ageing (Zhou et al. 2020) drawing attention to the effects of ageing on the neurodegenerative disorders. Kress and others elegantly demonstrated that not only glymphatic flow becomes dramatically reduced upon ageing (Kress et al. 2014), but CSF production and pressure decrease as well (May et al. 1990; Jessen et al. 2015). This decrease in CSF flow is postulated to be linked to increase in vascular stiffness and reduced brain artery pulsation (Kress et al. 2014; Mestre et al. 2017; Mestre, Hablitz, et al. 2018; Benveniste et al. 2019). Moreover, ageing was also associated with worse sleep quality (Mander, Winer, and Walker 2017), which in turn is linked to increased Aβ levels (Varga, Wohlleber, et al. 2016). In fact, sleep disruption is correlated to an increased Aβ deposition (Ju et al. 2013) and connected to cognitive dysfunction (Varga, Ducca, et al. 2016).

Noteworthy, during ageing not only glymphatic but also interconnected cerebral lymphatic network undergoes deterioration resulting in severely impaired drainage towards peripheral lymph nodes (Da Mesquita et al. 2021; Ma et al. 2017). Indicatively, studies performed in 5xFAD mice (an EOAD mouse model) showed that disruption of meningeal lymphatic vessels in younger animals (5-6 m) led to exacerbation of disease with observed increase in Aβ deposition and reduced extracellular clearance (Da Mesquita et al. 2018). More recently, the same authors demonstrated that in aged mice (13-14 m) deterioration of lymphatic vasculature was also observed (Da Mesquita et al. 2021). These 5xFAD mice rapidly develop amyloid pathology with plaque deposition starting around two months and spreading throughout cortex, hippocampus, thalamus, olfactory bulb and brainstem and even the spinal cord, but being absent in cerebellum. Although not assessed in our current studies, it would be interesting to explore also this glymphatic-lymphatic axis in Tet-Off APP mouse model using more sensitive tools, particularity in view that we did not observe significant difference in the outflow time constant τ_out_ between AD and control mice (Table 1&2). We conjecture that this might be related to the above discussed impact of ageing on vasculature, efficacy of (g)lymphatic system and CSF production in our middle-aged mice making either a difference in τ_out_ between both groups negligible and/or being beyond the detectability levels by MRI.

Furthermore, we speculate that changes in the glymphatic flow of AD mice may be in part related to astrocyte changes given that an extensive astrogliosis was observed, not only in the vicinity of amyloid-β deposits, but throughout the brain and also in regions devoid of lesions such as the brainstem (Figure 6C). Astrocytes are believed to be key players in the modulation of CSF-ISF exchange facilitated by AQP4 water channel at the astrocytic end-feet (Iliff et al. 2012; Mestre, Hablitz, et al. 2018; Hablitz et al. 2019; Harrison et al. 2020). Deletion of AQP4 in middle-aged APP/PS1 mice (12 m) increased amyloid-β accumulation, vascular amyloidosis and aggravated cognitive deficits (Xu et al. 2015). The perivascular localization and expression of AQP4, required for efficient CSF-ISF exchange, was shown to decline with age (Kress et al. 2014), and in AD patients AQP4 polarization was associated with disease stage (Zeppenfeld et al. 2017). In addition, the extent of vascular amyloidosis in EOAD murine models was closely correlated with astrocyte polarization (Yang et al. 2011; Kimbrough et al. 2015). In view of the aforementioned literature and the wide-spread astrogliosis in our AD mice with high levels of GFAP-positive astrocytes in regions of high Aβ burden, the changes in AQP4 polarization and coverage between both groups would be intuitively expected. However, our sample size and experimental outcome were not sufficient to draw currently reliable conclusions in this regard (Figure 6E). Moreover, astrogliosis is a process via which astrocytes react to different forms of neuropathology like Aβ insult when astrocyte functions are altered (Zlokovic 2011; Kimbrough et al. 2015). Thus, while healthy astrocytes play a vital role in neuro-glia-vascular unit by connecting the vasculature to neurons to ensure neuronal communication and energy demands (Iadecola and Nedergaard 2007), the astrogliosis contributes to pathology of AD by loss of astrocytes normal function and toxic effects (Zlokovic 2011). To this end, the loss of astrocyte polarization was suggested to be a consequence rather than a cause of Aβ-deposition with observed impairment of gliovascular coupling (Kimbrough et al. 2015). Additionally, we have demonstrated the effects of soluble Aβ on the resting-state neuronal networks at the pre-plaque stage and thus preceding the reactive astrogliosis observed in the presence of wide-spread amyloid plaque deposition (Ben-Nejma et al. 2019).

While it is still impossible to establish to what extent disruption in global brain activity and CSF circulation-drainage play a role in the initiation of AD pathology, we believe that a bidirectionally detrimental influence is the most likely scenario and should be carefully considered by combining different models and tools. This becomes particularly important in view of recent exciting studies that indicated to possible cognitive improvement by decreasing Aβ load either via combining halted APP overexpression and immunotherapy in Tet-Off APP mice (Chiang et al. 2018), or using targeted enhancement of (g)lymphatic CSF-lymphatic clearance in 5xFAD mice (Lee et al. 2020; Da Mesquita et al. 2021; Lin et al. 2021).

## CONCLUSIONS

To conclude, in our studies we chose to investigate Tet-Off APP mice and control littermates to be able to access and dissociate the effects of pathology and ageing on the global efficacy of CSF-ISF exchange in the context of neuronal network dysfunction, at a time point mimicking advanced stage LOAD. It would be very interesting to further extend these studies by choosing different mouse-age regimes for APP expression, in analogy to the work of Jankowsky and colleagues (Wang et al. 2011), and/or transgenic models overproducing Aβ with APP expressed at physiological levels. Given the interesting experimental outcome, we envision it will be highly valuable to perform future studies of the glymphatic-lymphatic axis in relation to network dysfunctions in neurodegenerative disorders both at the whole brain scale by MRI as well as at the microscale by immunohistochemistry and other techniques. Such studies will allow a closer look on the effects of amyloid beta on brain fluid circulation and the link to brain network dysfunctions.

## ABBREVIATIONS

Aβ: amyloid-beta
AD: Alzheimer’s disease
ANTs: Advanced Normalization Tools
APP: amyloid precursor protein AQP4: aquaporin-4
AUC: area under the curve
CamKIIα: calmodulin-dependent protein kinase type II alpha chain
CB: cerebellum
CM: cisterna magna
CSF: cerebrospinal fluid
CTL: control
DCE-MRI: dynamic contrast-enhanced magnetic resonance imaging
DMN: default mode (like) network
DOX: doxycycline
EOAD: early-onset of Alzheimer’s disease
Gd: gadolinium
Gd-DOTA: gadoteric acid
GS: glymphatic system
Gcv: cerebral vein of Galen
HC: hippocampus
ISF: interstitial fluid
LOAD: late-onset of Alzheimer’s disease
MREG: magnetic resonance encephalography
NONE: non-injected
NREM: non-rapid eye movement
NTg: non-transgene carrier
OB: olfactory bulb
OlfA: olfactory artery
PCA: principal component analysis
Rrhv: rostral rhinal vein
SAL: saline
Sss: superior sagittal sinus
Trs: transverse sinus
ROI: region-of-interest
rsfMRI: resting-state functional magnetic resonance imaging
tetO: tetracycline-responsive
tTA: tetracycline-Transactivator

## DECLARATIONS

**ETHICS APPROVAL**

All experiments were approved and performed in strict accordance with the European Directive 2010/63/EU on the protection of animals used for scientific purposes. The protocols were approved by the Committee on Animal Care and Use at the University of Antwerp, Belgium (permit number 2014–76).

## CONSENT FOR PUBLICATION

Not applicable

## AVAILABILITY OF DATA AND MATERIALS

Raw data and images are available upon reasonable request from the corresponding authors.

## AUTHORS’ CONTRIBUTIONS

IBN, AJK, AVdL and GAK conceived and designed the study. IBN performed the surgeries, MRI and optical images data acquisition. IBN, GAK analysed MRI data. GAK developed scripts and pipeline used for MRI data processing, analysis and supervised MRI data analysis. VV, IBN, AJK contributed to MRI sequence adjustment. VV and IBN did the MRI data pre-processing. AJK, IBN, PP developed protocols and contributed to ex-vivo immunostaining studies. AJK supervised all aspects of the daily work and performed data interpretation. IBN, AJK, GAK prepared figures and wrote the manuscript. All authors read, edited and approved the final version of the manuscript.

## ACKNOWLEDGEMENTS

The authors thank to Prof. Dr. Joanna L. Jankowsky (Baylor College of Medicine, Houston, Texas, United States) and Prof. Dr. JoAnne McLaurin (Sunnybrook Health Sciences Centre, Toronto, Canada) for providing single transgenic tTA and APP male mice. We also express our gratitude to Dr. Jelle Praet for initial help in establishing and managing mouse colonies. We are indebted to Jasmijn Daans for help in performing immunostaining experiments. We thank other members of Bioimaging lab, Johan Van Auderkerke for technical support and Prof. Marleen Verhoye for initial technical suggestions and discussion during meetings. The computational resources and services used in this work were provided by the HPC core facility CalcUA of the Universiteit Antwerpen, the VSC (Flemish Supercomputer Center), funded by the Hercules Foundation and the Flemish Government – department EWI.

## SUPPLEMENTARY DATA

### Voxel-based analysis reflects differences in the distribution of Gd-DOTA across groups

To assess the differences in the spatial distribution of the Gd-DOTA between the CTL and AD groups, voxel-based analysis (VBA) was performed at four time-points after injection (30, 60, 90, 120 min). To this end, a two-sample t-test was performed on a voxel-by-voxel basis comparing the percent signal change from baseline across the CTL and AD groups. No statistical significance was observed after multiple comparison correction, but Additional Figure 1 presents uncorrected statistics (p<0.05, uncorrected) that indicate a general agreement with our ROI-based and cluster-based analyses. Specifically, percent signal change was higher in the AD group in areas nearby the injection cite (pons, medulla; orange/yellow) as also observed in other analyses. In addition, VBA revealed signal decreases in AD relative to CTL in caudal areas close to the superior sagittal and transverse sinuses (blue) but note that signal intensity in those areas was low and within the variability observed in the non-injected groups and thus it is difficult to make robust conclusions.

**Additional Figure 1.**
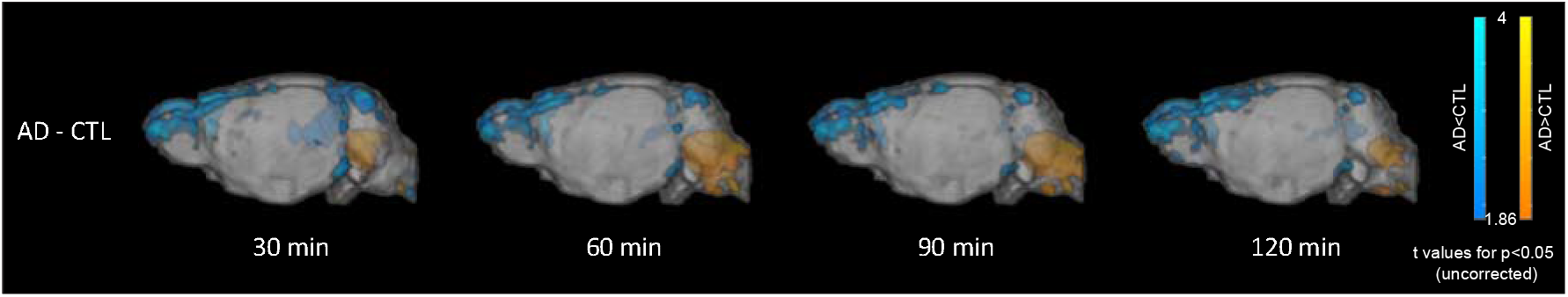
Voxel-based analysis (VBA) comparing signal intensity of CTL and AD mice at four time-points after injection (30, 60, 90, 120 min). The AD group demonstrated stronger signal intensity in regions adjacent to the infusion spot (orange/yellow) in all time points and decreased signal intensity in areas proximal to the superior sagittal and transverse sinuses (blue). The colour scales indicate T-statistic values, with orange/yellow representing the voxels significantly higher in the AD group and blue/cyan representing the voxels significantly higher in the CTL group (p<0.05, uncorrected).

